# Spatial position relative to group members affects weight gain in meerkats (*Suricata suricatta*)

**DOI:** 10.1101/2024.11.29.625976

**Authors:** Rasekuwane Mosia, Vlad Demartsev, Aliza le Roux, Marta B Manser, Ariana Strandburg-Peshkin, Lily Johnson-Ulrich

## Abstract

Social animals often face a trade-off between the costs of foraging competition among group members and the benefits of protection from predators offered by group living. The spatial position of an individual in relation to the other group members during foraging can mediate the effects of this trade-off as individuals at the front or edge may have better access to food resources, but also higher predation risk than individuals near the centre of the group. Using meerkats (Suricata suricatta) as a model species, we investigated the effect of individual spatial position within a group on foraging success. We determined the spatial position of individuals in a meerkat group by fitting the animals with high-resolution GPS loggers. As a proxy of foraging success, we used meerkats’ individual body weight differences between the start and the end of daily data collection over foraging periods (3 hours). We found significant individual differences in meerkats’ spatial positions within the group. In addition, age-dependent differences in spatial position became obvious, with younger individuals spending more time in the centre of the group and less time in the front. However, younger individuals who spent more time on the side of the group relative to older individuals had higher daily weight gain, indicating more successful foraging. In older individuals, we found that the dominant females tended to spend more time towards the front of the group, but gained less weight in this location, contrary to the predicted association between front edge of the group and better access to food resources. Our results suggest that the relationship between weight gain and spatial position is highly nuanced and likely dependent on more than just trade-offs between foraging success and predation risk.

**Highlights:**

- Spatial position relative to other group members during foraging was a highly repeatable trait for individual meerkats, though time spent near the front of the group was more strongly influenced by individual traits (age, sex, and social rank)
- Younger meerkats spent less time in the front and more time in the centre of the group relative to older meerkats, but had the highest foraging success towards the sides of the group
- Dominant female meerkats spent more time near the front of the group, but had reduced foraging success in this position
- Meerkats may trade-off more than just foraging success and predation risk when making decisions about where to position themselves relative to other group members

## INTRODUCTION

Collective movement of animal groups has been described across taxa including herds of mammals, swarms of insects, schools of fish, and flocks of birds (Bode et al., 2010; Farine et al., 2014; Giardina, 2008). The advantages of moving collectively largely overlap with the advantages of social living. In many species, individuals benefit from moving with group members because of increased anti-predator vigilance and higher success in between-group competition for food resources (Bednekoff & Lima, 1998; Lehmann et al., 2016). For example, blue monkeys (*Cercopithecus mitis*) are more vigilant when in lower foliage density which is associated with higher predation risk, but reduced their vigilance in the presence of kin with which they form strong affiliative relationships with (Gaynor & Cords, 2012). Further, animals that move collectively can benefit from the sharing of information about the environment. For example, schools of golden shiners (*Notemigonus crysoleucas*) were able to track environmental gradients better when moving in larger groups (Berdahl et al., 2013). However, collective living also comes with challenges, such as coordinating optimal decisions among all group members (Conradt & Roper, 2005; Petit & Bon, 2010) and facing intra-group competition over resources such as food (Arseneau-Robar et al., 2023; Holekamp & Sawdy, 2019).

When moving collectively animals may mitigate some of the trade-offs between competition and predation risk by occupying different relative spatial positions within the group, which may vary in their predation risk and access to resources (Hall & Fedigan, 1997; Hirsch, 2007; Tkaczynski et al., 2014). While individuals positioned at the front or at the edge of the group might be subjected to higher predation risk, they may also gain better access to food resources compared to those at the centre or the back of the group (Rayor & Uetz, 1990; Rowcliffe et al., 2004; Stahl et al., 2001). For instance, foraging success increases towards the edge of the group in fallow deer (*Dama dama)* (Focardi & Pecchioli, 2005). These cost-benefit trade-offs may also be influenced by dominance hierarchies as dominant individuals are often found in locations that are associated with lower costs and higher benefits (Hall & Fedigan, 1997; Murray et al., 2007). In particular, when food resources are scarce, individuals that are located towards the edge of the group may have increased foraging success or food intake (Morrell & Romey, 2008). However, the edge positions are not necessarily the preferred locations for all group members, as dominant individuals in the centre of the group can monopolize food patches or employ scrounger tactics to improve their foraging success (Grant et al., 2002; Hirsch, 2007; Murray et al., 2007).

Quantifying foraging success directly is challenging, and most research on these trade-offs uses indirect methods of estimating foraging success such as counting the number of feeding events per minute (Hintz & Lonzarich, 2018). Here, we aimed to investigate the trade-off between spatial position and foraging success, estimated by individual weight gain during the foraging period in meerkats (*Suricata suricatta*). We used weight gain as a measure of foraging success because it takes into account the cumulative effect of numerous foraging events over time and, importantly, *the quantity of food acquired*, which other indirect measures of foraging success may fail to account for.

Meerkats live in groups of up to 50 individuals, and they are a cooperatively breeding species living in arid parts of Southern Africa (Clutton-Brock & Manser, 2016). During the day, they maintain a cohesive group structure and forage together on dispersed subterranean prey while moving through their territory (Doolan & Macdonald, 1996). Meerkat groups consist of a dominant pair that monopolizes breeding and subordinate individuals, mostly offspring of the dominant pair, that help raise the young (A. F. Russell et al., 2003). Previous research has found that dominant meerkats were frequently located towards the front of the group, presumably due to higher foraging success in the front (Averly et al., 2022; Gall & Manser, 2018). However, the link between spatial position and foraging success has never been tested in meerkats. Here, we investigated the potential link between spatial position and foraging success during cooler and drier winter months (June – September 2023), when prey availability is lower (Doolan & Macdonald, 1996), as any trade-offs between predation risk and foraging success are likely to be most apparent.

We predicted that individuals at the front of the group, which potentially encounter food resources first, would have increased weight gain at the end of each foraging session. We also predicted that, since the front position might be associated with better assessment of the resource distribution, (Focardi & Pecchioli, 2005; Gall & Manser, 2018) more experienced older or higher-ranking individuals would occupy this location more frequently. In addition, we expected younger individual meerkats, which are potentially more at risk from predation because they are less experienced in responding to predatory threats (Hollén et al., 2008), would spend more time in the centre or the back of the group where they might benefit from the collective vigilance of the group.

## METHODS

### Study site and population

We conducted the study at the Kalahari Research Centre (KRC), Kuruman River Reserve (28°58′S, 21°49′E), in the Northern Cape, South Africa. The meerkat population on site has been studied since 1993. Meerkats are habituated to human presence and marked with unique dye-marks making them easily identifiable at the individual level (Manser, 2018). Individual weights were routinely collected, up to three times per day on three to five days per week, by luring meerkats onto electronic scales using a small piece of hardboiled egg or a few drops of water (Manser, 2018).

### Ethical note

The study was approved by the University of Pretoria Ethical Committee (EC031-1, NAS003/2022) and the Northern Cape Department of Environment (NAS033/2022, EC031-17, EC047-16) and Northern Cape Province Department of Environment and Nature Conservation (FAUNA 1020/2016, FAUNA 0996/2022). We certify that our study complies with all applicable institutional norms as well as the ASAB/ABS guidelines for the use of animals in research and with the laws of South Africa, the country in which the study was conducted. All field procedures including collaring, focal follows, and weight collection are based on previously established protocols. Because meerkats are highly habituated to humans, weighing scales, and accepting water from water bottles we were able to both measure neck sizes and deploy custom-fitted collars without the need for anesthetization while meerkats took a few sips from a water bottle (Video 1A; also see Methods: Collar deployment). While most meerkats showed no reaction to the attachment of a collar, some meerkats showed momentary confusion (e.g., rolling in the sand immediately after a collar was attached). However, all meerkats resumed their previous behaviour less than one minute after deploying the collar. We observed no behavioural changes in meerkats as a result of wearing collars or changes in their habituation towards human observers. If meerkats showed any signs of ongoing discomfort (e.g., persistent scratching or rolling) or if we observed the collar was fitted too loosely it was removed immediately (N ∼ 1 and 4 instances respectively out of 71 collared meerkats). All collars were removed within 24 hours of the end of the data collection period (Video 1B). For detailed information on collar deployment see the Supplementary Information from Averly et al. (2022) which followed an identical protocol. Previous research on the effects of radio collaring meerkats have likewise found no effect of collars on predation rates or foraging efficiency of (Golabek et al., 2008).

**Video 1.**
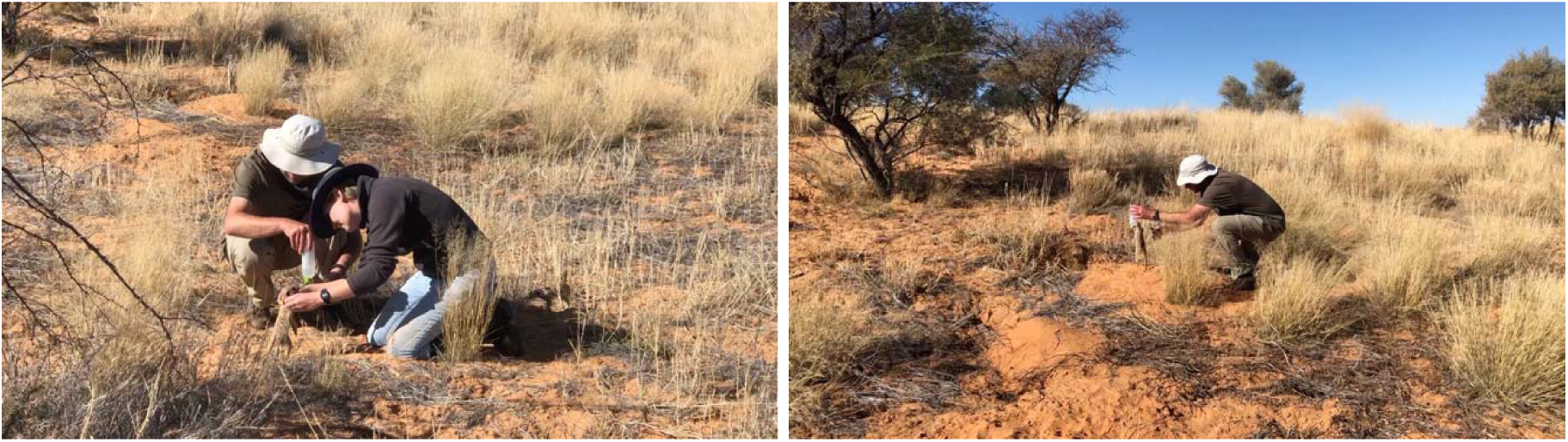
A) Left: VD and LJU deployed a collar while the meerkat drinks from a water bottle. B) Right: VD snipping the leather strap of a collar to remove it while the meerkat drinks from a water bottle. Observers always washed their hands and/or used hand sanitizer before and after close contact with meerkats.

### Long-term data and individual traits

All meerkat groups in this study were part of detailed longitudinal monitoring that includes daily records over three to five days per week of behaviour, group composition, pregnancy, disease, injuries, and life history data. We used a binary variable for age that merged several age categories traditionally used in studies with meerkats: individuals that were older than 1 year were considered ‘older’ individuals and this category included dominant females, dominant males, yearlings (1 to 2 years), and adults (2 years and older). Individuals younger than 1 year were considered ‘younger’ individuals and this category included subadults and juveniles (see Table 1 for detailed group composition and age classification). We chose 1 year of age as a binary cutoff because this roughly represents the age of sexual maturity in meerkats (English et al., 2013). All meerkats were assigned a date of birth based on physical signs of parturition in the mother, e.g, a sudden loss in weight and signs of lactation. The age categories of juvenile, subadult, yearling, and adult are based on the typical ages of life-history milestones (e.g., foraging independence, sexual maturity, dispersal). Individuals are sexed as pups based on the ano-genital distance and sexes are further confirmed by the later appearance of sexual characteristics (e.g., the appearance of the scrotum). Dominance is determined based on aggressive and submissive interactions with other same-sex group members alongside behavioural changes (e.g., anal marking) or physiological changes (weight) (Duncan, 2021).

**Table 1.**
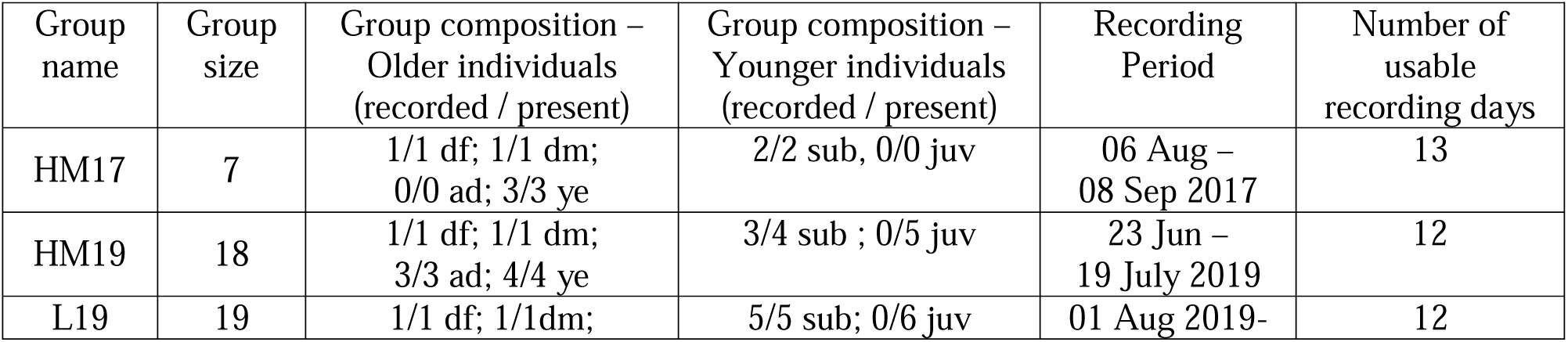
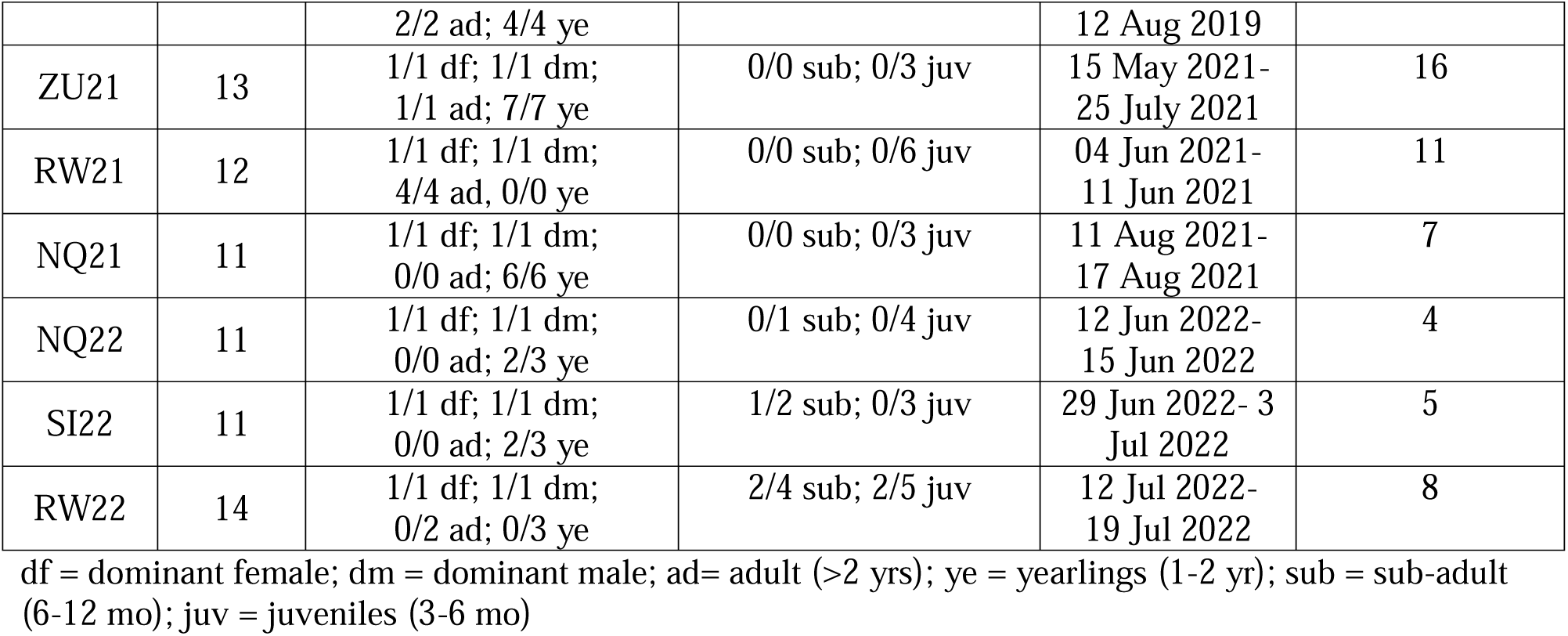
Summary of group composition and data collection.

### Data collection periods

We obtained data from a series of short-term deployments of meerkat groups between August 2017 and July 2022 (Table 1). The deployments covered nine different groups in total.

### Spatial positioning: Collar deployment

We deployed custom-built collars on as many individuals in a group as it was possible, without the need for anesthetization. We lured meerkats to stand up bipedally using a water bottle, then secured the collars using a custom designed magnetic clasp. The closing system of the collar included two magnets (measuring 1 x 5 x 5 mm) affixed to 3D-printed plastic clasps positioned at both ends of the leather band. This design aimed for effortless closure while necessitating human action to open it.

We constructed each collar according to the neck size and weight of the individual meerkats, with the collar weight (22-26 g) not exceeding 5% of the animals’ body weight, in accordance with Golabek et al. (2008). Attached to the collar was a GPS tag (Gipsy 5 in 2017 and 2019, Axy Trek Mini in 2021; Technosmart, Colleverde, Italy) that recorded the animals’ coordinates at 1 fix per second. Only meerkats over 500 g (i.e., most meerkats other than juveniles) were collared with GPS tags. We programmed all units to activate daily, for three consecutive hours, during times when meerkats engaged in group foraging. At the end of data collection, we removed the collars by snipping the leather straps with a mini diagonal cutter.

### Spatial positioning: Focal recording

For individuals that were already equipped with a long-term radio collar or could not otherwise be collared, we performed focal data collection by following the individual with a telescopic pole with a GPS tag attached to it. Here, the observer kept within 1 m of the foraging meerkat for the 3-hour duration of each daily session. At the same time, the observer recorded voice notes of the focal meerkat’s behaviour (e.g., eating, calling, and grooming) including noting occasional moments when the meerkat moved more than 2 m away from the GPS tag (these time segments were then removed from the recorded trajectories).

### Spatial categories calculation

From the recorded individual GPS coordinates we calculated the group ‘centroid’ by averaging all individuals’ GPS locations each second. We used a 10 m spatial discretization to calculate the heading of the group based on previously established methodology (Averly et al., 2022). Briefly, the heading at a given time was defined as the vector pointing from the centroid’s past location after it had moved a distance of 10 m to its present location at the given time. We then calculated each individual’s location relative to the group centroid and heading at every second. We then summarised information about spatial position over the three-hour data-collection session using three different approaches:

1: Relative Location: We first classified individuals as either ‘centre’ or ‘edge’ based on whether they were closer or farther than the median meerkat’s distance to the centroid at each second, such that half of meerkats in the group were categorized as centre and half as edge each second. Within the edge category, we classified individuals that were in the front quarter (between -π/4 and π/4 radians) as ‘front’. We classified those that were behind the centroid (greater than ± 3π/4 radians) as ‘back’, and those that were between π/4 and 3π/4 or -π/4 and -3π/4 as ‘side’. We then calculated the proportion of time each meerkat spent in these four locations (front, side, centre, and back) in each of the three-hour daily data collection sessions. We developed this measure of position in order to distinguish both entre from edge positions and within edge, to distinguish front, back and side positions in line with other research on spatial positioning outside of meerkats (e.g., Focardi & Pecchioli, 2005; Morrell & Romey, 2008).

2: Binary Location: We calculated the proportion of time spent in front half (binary position) of the group to replicate results from Averly et al. (2022).

3: Ranked Location: We ranked individual meerkats based on their front-back distance at each second and took the mean rank over the three-hour data collection session (‘ranked position’) to replicate results from Gall & Manser (2018). Rank was scaled from -1 to 1 where a rank of 1 indicated that a meerkat was the furthest towards the front of the group and a rank of -1 indicated that a meerkat was the furthest towards the back of the group. Rank thus reveals which individuals were, on average, far ahead or far behind other group members across a continuous scale whereas relative location and binary location are categorical.

### Foraging success: Change in body weight gain

To determine individual daily weight change, we followed the standard weighing procedure at the study site and weighed meerkats every morning, before they began foraging, and at the end of the daily three-hour data collection session. We used the change in body weight (Δweight) between the two daily weighing sessions as a proxy for individual foraging success. To control for average weight of meerkats of different age and sex classes, we standardized each individual’s daily Δweight (calculated z-score of Δweight) against the average Δweight values, which was determined by calculating the rolling mean for individual weight over 11 weight sessions; +/- five sessions including the current one (Δweight-Z). To control for the daily variation in the foraging success of the group due to unobserved environmental factors (e.g. weather), we also standardized individual daily Δweight-Z against the average Δweight-Z of meerkats in the same group on the same day (Δweight-Z2). The z-scores for both Δweight-Z and Δweight-Z2 were calculated using the formula: z-score = (x-μ)/σ where μ and σ are the average and standard deviation for an individual meerkat’s Δweight over 11 weights for calculating Δweight-Z. For Δweight-Z2, μ and σ are the average and standard deviation of Δweight-Z over a single day for the entire group.

### Statistical Analysis

To test our predictions, we first determined how consistently individual meerkats occupy certain spatial locations while foraging. We tested if the spatial preferences were affected by age, sex, dominance rank, and individual identity. Second, we also investigated whether weight gain was affected by the proportion of time spent in different locations and whether the effect of spatial location on weight gain is also affected by age, sex, or dominance rank.

We created generalized linear mixed models with the R package ‘glmmTMB’ (Brooks, Mollie et al., 2017; R Core Team, 2023), to determine whether individuals showed consistency in within-group spatial positioning. In these models we used position as the response variable and age (older vs younger individuals), dominance rank (dominant vs subordinate), and sex (female vs male) as predictor variables with a random effect of individual ID. Since in meerkats the dominant female fills a critically different social role from the dominant male (Averly et al., 2022), we also included a rank by sex interaction. We calculated R values for individual ID using the R package ‘rptR’ (Stoffel et al., 2017) for models with and without fixed effects.

Next, we investigated whether within-group location predicted weight gain. In these models we used both measures of standardized weight gain as the response variables. Other factors such as sex (male or female), rank (dominants or subordinates), age (older individuals or younger individuals), and a random effect of individual ID were also included so that their effects could be controlled for. All additional variables were included as moderators of the location-weight gain relationships (i.e., they were included as interactions with location). We also included an interaction between rank and sex for the same reasons specified above. We used the R packages ‘emmeans’ and ‘modelbased’ to estimate marginal means and marginal slopes from models (Lenth, 2022; Makowski et al., 2020). For marginal means and slopes we focus on four dominance-sex-age categories, dominant female, dominant male, subordinate older individuals and subordinate younger individuals because we only expected (and only found) sex differences among the dominant individuals. We include estimates for all six possible combinations of these factors in the Appendix (Figure A1).

## RESULTS

### The effect of age, sex, and dominance on within-group spatial position

Using ‘relative location’ (front, back, side, or centre), we found that most individuals spent roughly half of their time in the centre of the group (Figure 1) which was expected from our definition of the group centre as the median meerkat distance from the group’s centroid. However, relative to older individuals, younger individuals spent significantly less time in the front (Table A1, estimate ± SE= -0.024 ± 0.011, p= 0.049) and slightly more time in the centre (Table A1, estimate ± SE= 0.078 ± 0.027, p= 0.004). We also found highly repeatable effects of individual ID for back, side, and centre locations (Table A1, R = 0.11 – 0.24, p < 0.05) regardless of the inclusion or exclusion of fixed effects. These results suggest that individuals tend to spend a consistent amount of time in these relative locations (back, side, and centre) that is not explained by age, sex, or rank. While we did find repeatable effects of individual ID for the front location, this was only the case before controlling for the effect of age, sex, and rank (Table A1, R= 0.131, p = 0.008).

**Figure 1.**
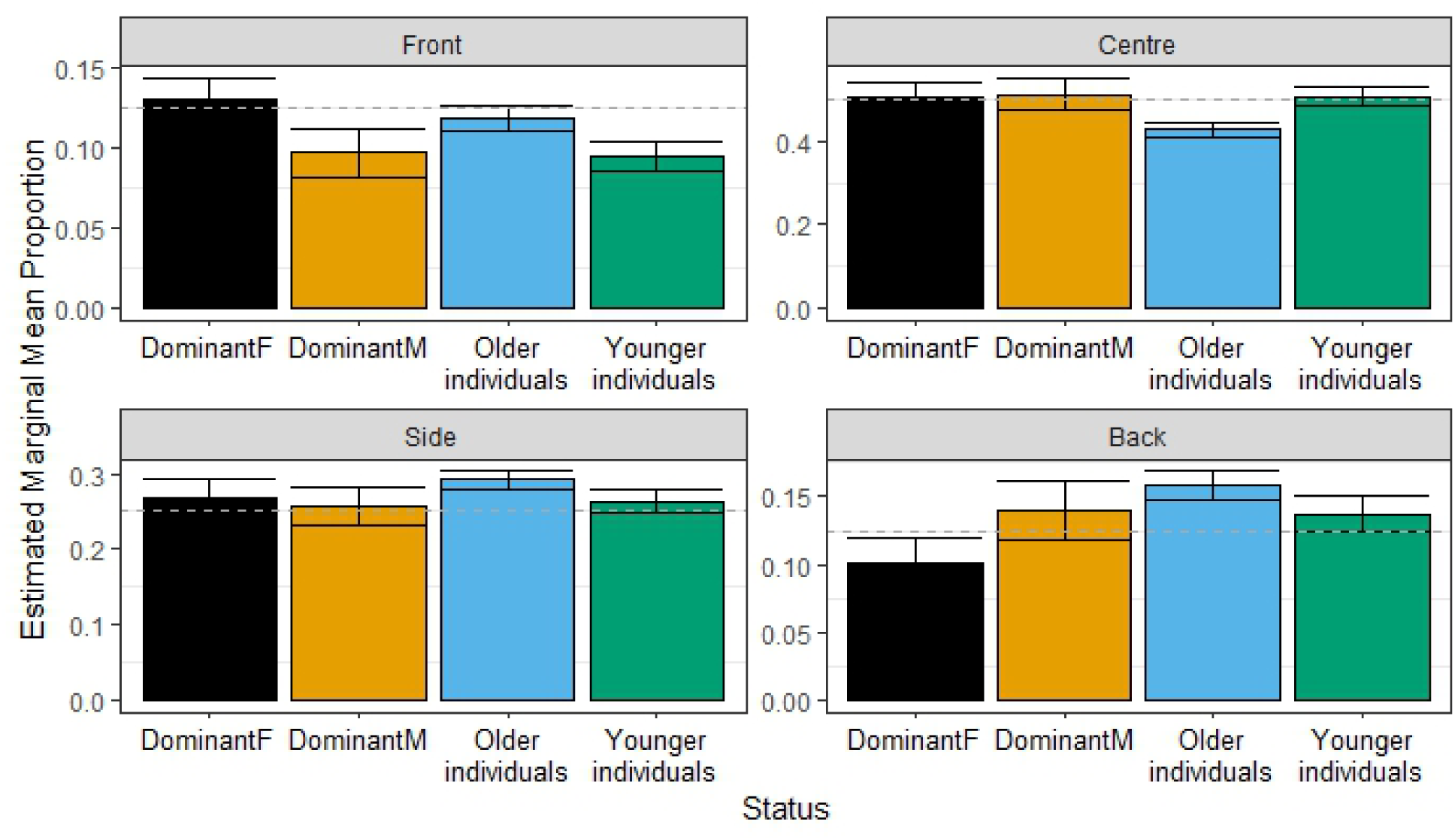
The proportion of time (given in %) spent in different spatial positions by meerkats within the group. The consistency of spending time in different locations is measured using the social status of individuals to control for its effects. The emmeans is the marginal means of the proportion of time spent by individuals in different locations (from model results reported in Table A1). The letters F and M at the end of ‘Dominant’ are for sex (Female and Male). The horizontal dotted line indicates the proportion of time expected by chance in each location.

The effect of age-rank-sex status on binary location (front vs back) revealed a significant interaction between rank and sex. This interaction appeared to be driven largely by the effect of dominant females, which spent significantly more than 50% of their time in the front-half of the group (95% CI: 0.539-0.646, p= 0.008). However, post-hoc pairwise comparisons between age-rank-sex classes were non-significant (p > 0.05 for all contrasts after Tukey adjustment). There was also significant, repeatable individual variation in the amount of time meerkats spent at the front or at the back of the group before controlling for the effect of age, sex, and rank (Table A2, R= 0.117, p = 0.013) but not after, which suggests that the effect of individual ID on binary position can be mostly explained by the effects of age, sex, and rank.

In the ranked location model, the interaction between rank and sex was not significant and there were no significant post-hoc pairwise comparisons between rank-sex classes. While our model output showed a significant effect of rank after controlling for the interaction between rank and sex (Table A2), the total effect of rank was not significant (post-hoc estimated marginal mean for dominant vs subordinate: E= 0.06, p = 0.18). There was also significant individual variation in ranked location (mean front-back distance) only before controlling for the effect of age, sex, and rank (Table A2, R= 0.106, p = 0.024), again suggesting that any individual variation can be explained by variation in age, sex, and rank.

### Spatial location and weight gain

For meerkats’ relative location (front, back, centre, and side), side had a significant effect on Δweight-Z for younger meerkats (Figure 2). Younger individuals that spent slightly more time on the side relative to older individuals significantly gained more weight (Table A3, E = 4.10, p = 0.011), but this was only marginally significant for Δweight-Z2 (Table A5, E = 2.61, p= 0.087). Whereas Δweight-Z was standardized against an individual’s average weight gain, Δweight-Z2 was standardized against both an individual’s average weight gain and the average Δweight-Z for the group on a particular day. Meerkats’ binary location had a significant effect on Δweight-Z and Δweight-Z2 for dominant females, which gained less weight in the front-half of the group (Table A4 & A6, E= -2.82 – -2.89, p= 0.025). In addition, subordinate older individuals gained significantly more weight in the front-half of the group, but only for Δweight-Z2 (Table A6, E = 1.89, p = 0.026). Ranked position had a significant effect on weight gain only for dominant females, who gained less weight when spending more time further ahead of the group (Figure 3; Table A4, E = -1.78, p = 0.02; Table A6, E = -1.68, p = 0.02).

**Figure 2.**
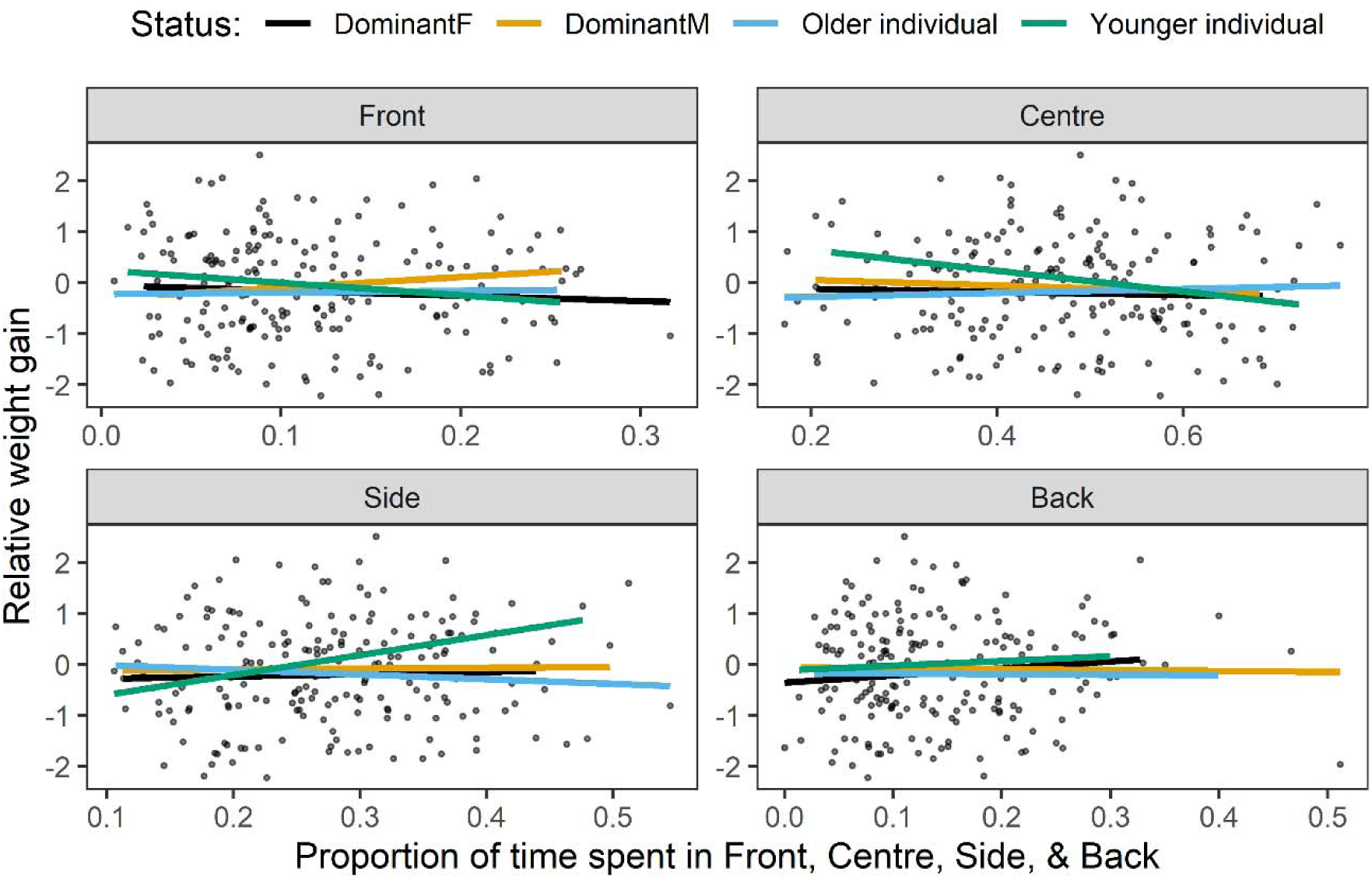
The effect of individuals’ spatial position on Δweight-Z (standardized to mean individual weight change) including the effect of age and social rank. The x-axis is the proportion of time spent in different positions each day. The y-axis is the standardized weight gain. The lines show the best fit.

**Figure 3.**
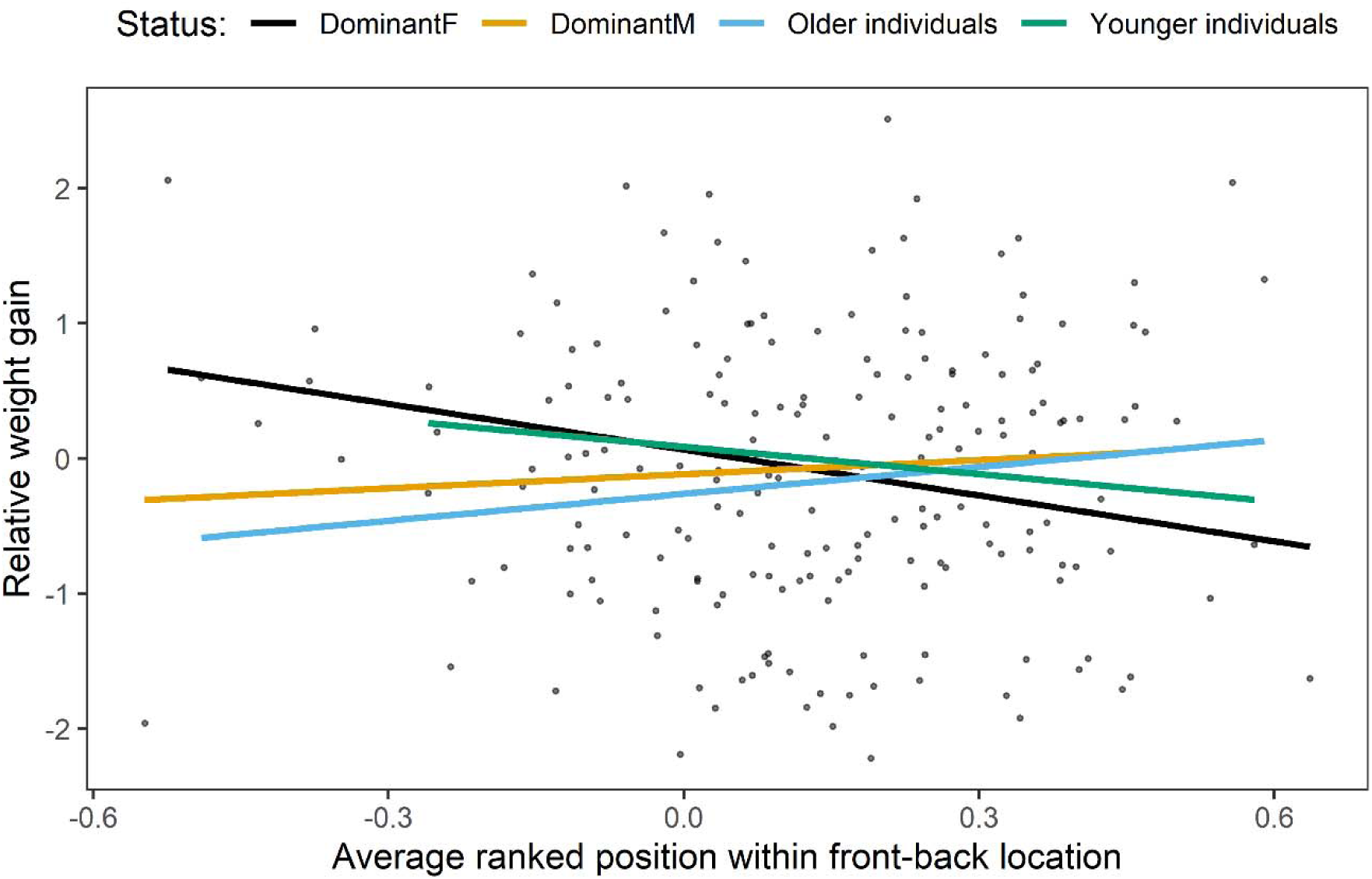
The effect of ranked front-back location on Δweight-Z (standardized to mean individual weight change) on relative weight gain of different individuals according to their social status. Younger individuals and adults refer to subordinates. The x-axis shows the average rank per foraging session on a scale of -1 to 1 where a rank of 1 indicates the meerkat occupied the position furthest ahead and a rank of -1 indicates the meerkat occupied the position furthest behind. A rank of 0 indicates the meerkat occupied the median position along the front-back axis. The y-axis is the standardized weight gain. The lines show the best fit.

## DISCUSSION

We examined relative spatial position in foraging meerkats applying three models and found that meerkats of different age classes and dominance ranks consistently chose specific locations within the group. Younger individuals were consistently more often in the centre of the group compared to older individuals when using the relative location metric (front, back, side and centre) model. An interaction between sex and dominance rank revealed that dominant females spent more time in the front location, and this was confirmed when applying a binary location (front vs back) model. In contrast to the expectation that individuals at the front may gain more weight, we found that younger individuals that were on the side edges had higher daily weight gain relative to other age classes, suggesting higher foraging success for them in this location. Also, in contrast to our expectations, dominant females gained less weight when spending more time towards the front of the group in both our binary location model and ranked location model (front to back on a scale of 1 to -1).

We found significant individual repeatability for the proportion of time meerkats spent in back, side, and centre relative locations. However, for the tendency to occupy the front relative location, front binary, and front-back ranked location, our results showed that any effect of individual ID is largely superseded by age, sex, and rank. These results suggest that frontness may be more dependent on age, sex, and rank whereas the relative amount of time spent towards the side or centre of the group may depend on other factors related to individual identity. Indeed, meerkats’ choice of spatial position while foraging may be dependent on the complex interaction of several factors including opportunities for information-transfer (Goodale et al., 2010), perception of safety (Hirsch & Morrell, 2011; Janson, 1990), familiarity with the area (Wolf et al., 2009), optimal foraging theory (Davis et al., 2022), leadership (Averly et al., 2022), and social status in the group (Stahl et al., 2001). Given our significant repeatability results, it appears that individual location in the group is not random but likely governed by socio-ecological factors that affect individual decision-making about relative spatial position.

Juvenile animals are often targeted by predators (Barber-Meyer & Mech, 2008), and predation pressure often increases towards the edge of the group (Krause, 1994). We suspect that younger subordinate meerkats position themselves in the centre of the group for relative safety at the expense of foraging success which is higher for them on the side edge of the group. Predation avoidance has been shown to drive juvenile individuals towards the centre of the group in white-nosed coatis (*Nasua narica*) (J. K. Russell, 1980) and to lower juvenile brown capuchin monkeys’ (*Cebus apella*) foraging effort (Janson, 1990) which also had more foraging events at the edge of the group (Di Bitetti & Janson, 2001). The sides of the groups, similarly to the front edge, might be less exploited in comparison to the centre and the back. Additionally, side edges might be safer than front/back, thus offering younger meerkats a balance between predation risk but a more rewarding foraging environment. Alternatively, younger meerkats might avoid competition with conspecifics in the denser centre of the group. Similarly, as the front of the group is often occupied by dominant and otherwise influential older meerkats, younger subordinate meerkats might be unable to efficiently exploit the available resources there. Similar findings were also shown for subordinate males in cichlid fish, *Astatotilapia burtoni,* who spent more time in the periphery (edge) of the group because aggressive, dominant males occupied central locations (Rodriguez-Santiago et al., 2020). Juvenile ring-tailed coatis (*Nasua nasua*) also spent more time in the front edge of the group where predation is likely higher (Hirsch, 2011). These discrepancies are likely driven by the socio-ecological differences between the species determining the cost-benefit balance for occupying certain locations.

Dominant meerkats were found to be located more often towards the front of the group than in the other relative locations in several previous studies (Averly et al., 2022; Barnard, 2000; Bousquet et al., 2011; Gall & Manser, 2018). These studies analysed meerkat location as either binary (front vs back half) or a front-to-back ranked location but did not consider centre vs edge effects as we did here. When determining meerkat location using binary or ranked locations, we replicated the results of dominant females spending more time at the front-half. However, in contrast to our expectations and the predictions made in previous studies, we found that dominant females gained significantly less weight the more time they spent towards the front of the group for binary (for both Δweight-Z and Δweight-Z2) as well as ranked (Δweight-Z) locations. This suggests that dominant females likely do not spend more time at the front of group as a result of higher foraging success in this location. Instead, dominant individuals may spend more time towards the front because it is associated with stronger influence on the group heading and speed (Averly et al., 2022). However, dominant females tend to have high influence independent from the effect of frontness on influence (Averly et al., 2022). And, in our comparison considering more detailed spatial categories (centre, front, side, back), we found that while the dominant females were spending more time at the front-half of the group, they were not necessarily occupying the front edge of the group as measured by our relative position metric. Future research investigating the effects of relative spatial position on leadership, influence, foraging success, and predation risk from moment-to-moment, rather than aggregated over a morning foraging session, could reveal more about when and where meerkats are foraging, leading, or doing other behaviours.

These results also contrast with research in other species which finds that dominant individuals benefit most from spending more time in the centre when food items are slowly depleting (Hirsch, 2007) and reports that dominant capuchin monkeys were predominantly located in the centre (Hall & Fedigan, 1997). We also found that subordinate adults gained more weight when they spent more time in the front-half of the group in one of our models, but this finding was not consistent across other models.

Our study revealed novel results from distinguishing between foraging success not just between the front and the back of the group, but also the centre vs the edge of the group. Future work could investigate whether the spatial location method used in this study would reveal similar results in other collectively foraging animals. However, using weight gain as a measure of foraging success over a three-hour period fails to elucidate precisely where individuals were located when they had more successful foraging events. Our data were collected while meerkat groups were foraging, but individual meerkats can engage in other behaviours during these periods. Additionally, when reaching satiation after a successful foraging bout, individuals might change their relative location to one perceived as less risky or simply divide their time between the different locations evenly (CluttonLBrock et al., 1999). To further understand the effect of spatial position on foraging success and fitness in social animals, future studies will benefit from methods of detecting food intake events in combination with high-resolution GPS data and weight gain. These studies could capture the moment-to-moment location of foraging events rather than looking at relatively longer sections of time, allowing a more direct link between within-group positioning and foraging success to be drawn.

Finally, while we hypothesized that predation pressure is highest towards the front and edge of the group based on research in other animals (Hirsch & Morrell, 2011; Morrell & Romey, 2008; Rayor & Uetz, 1990; Tkaczynski et al., 2014), we observed no predation events during our data collection periods and therefore it remains unclear whether predation risk has an impact on where meerkats spend their time within the group. Furthermore, many of the meerkats’ predators are aerial and here the location in the group might be less important than for terrestrial predators. To further test hypotheses about the trade-offs related to spatial positioning, more data is needed on the actual or perceived costs of variation in location for meerkats. Ultimately, while we found several effects of location and age-rank-sex classes on weight gain, a great deal of variation in weight gain remained unexplained (model residual variance range = 0.84 – 0.89). This suggests that spatial position is likely to be one of many other factors that influence foraging success from day to day. It will be necessary for future studies to take into consideration the effect of seasons and habitats in which the study species are found. Similarly, the physical characteristics of environments, affecting movement and potentially shifting the costs of foraging effort must be considered.

## Conclusion

In sum, we found significant individual differences in the proportion of time spent in different relative spatial positions within a group, primarily due to the effects of age, sex, and rank. We also found effects of spatial positioning on weight gain for dominant females and younger meerkats, but a large amount of variation in weight gain was unexplained. Future work may benefit from linking our methods to more time-sensitive measures of foraging success, such as individual foraging events. In addition, future studies might also consider the types of food resources utilized by animals which may vary in their nutritional content. Our results highlight the power of combining high-resolution movement data to gauge spatial positioning with information on foraging success, as well as the benefits of investigating multiple measures of spatial position on foraging success.

## Author contributions

Conceptualization: M.B.M., A.S.P., V.D. and L.J.U.; Data curation: V.D. and L.J.U.; Formal analysis: V.D., R.M. and L.J.U.; Funding acquisition: M.B.M., A.S.P., R.M., Investigation: R.M., V.D. and L.J.U.; Methodology: A.S.P., V.D. and L.J.U.; Project administration: M.B.M. and A.S.P.; Resources: M.B.M. and A.S.P.; Supervision: A.l.R., V.D. and L.J.U.; Visualization: R.M and L.J.U.; Writing – original draft: R.M., V.D. and L.J.U.; Writing - review & editing: R.M., M.B.M., A.S.P., A.l.R., V.D. and L.J.U.

## Acknowledgments

We are grateful to T. Clutton-Brock, the Kalahari Research Trust and Northern Cape Department of Environment and Nature Conversation for research permission at the Kalahari Research Centre. We also thank T. Clutton-Brock for assistance and access to the weight data used in the present study. We thank the Universities of Zurich, Cambridge, Pretoria, and the MAVA foundation for supporting the field site. We thank T. Vink and W. Jubber for organizing the field site, the managers, and volunteers of the Kalahari Meerkat Project (KMP) for maintaining habituation and long-term data collection of the meerkats. The authors are also grateful to the surrounding farmers for allowing them to use their land we would like to thank all the members of the CCAS research group for their valuable input and advice on this study. We also thank Gabriella Gall (2017), Baptiste Averly (2019), Rebecca Schaefer (2019), Pauline Toni (2021), and Camille Lysemna (2021) for helping with data collection.

## Data Availability

R code and datasets are available at https://github.com/l-johnsonulrich/spatialForagingRepo.git

## APPENDIX

**Table A1.** The consistency of individual ID for the proportion of time spent in each position, including level of significance. Individuals’ consistency is explained by the effect of age, rank, and sex. All R values (R total ad R with fixed effects) are for the random effect of individual meerkats.

| Model | Predictor | Estimate | SE | P |
| --- | --- | --- | --- | --- |
| Front | Rank: Subordinate | -0.027 | 0.016 | 0.096 |
| Front | Age: Younger individuals | -0.024 | 0.011 | **0.049** |
| Front | Sex: Male | -0.033 | 0.019 | 0.077 |
| Front | Rank*Sex: Subordinate, Male | 0.022 | 0.022 | 0.066 |
| Front | R total (before controlling for fixed effects) | 0.131 | 0.078 | **0.008** |
| Front | R with fixed effects | 0.09 | 0.07 | 0.168 |
| Back | Rank: Subordinate | 0.042 | 0.023 | 0.069 |
| Back | Age: Younger individuals | -0.021 | 0.016 | 0.191 |
| Back | Sex: Male | 0.039 | 0.026 | 0.138 |
| Back | Rank*Sex: Subordinate, Male | -0.027 | 0.031 | 0.382 |
| Back | R total (before controlling for fixed effects) | 0.112 | 0.068 | **0.021** |
| Back | R w/ fixed effects | 0.11 | 0.068 | **0.015** |
| Centre | Rank: Subordinate | -0.031 | 0.031 | 0.321 |
| Centre | Age: Younger individuals | 0.078 | 0.027 | **0.004** |
| Centre | Sex: Male | 0.007 | 0.048 | 0.890 |
| Centre | Rank*Sex: Subordinate, Male | -0.004 | 0.056 | 0.921 |
| Centre | R total (before controlling for fixed effects) | 0.266 | 0.091 | **0.0002** |
| Centre | R w/ fixed effects | 0.243 | 0.078 | **4.35e-05** |
| Side | Rank: Subordinate | 0.011 | 0.027 | 0.468 |
| Side | Age: Younger individuals | -0.029 | 0.0184 | 0.114 |
| Side | Sex: Male | -0.011 | 0.032 | 0.724 |
| Side | Rank*Sex: Subordinate, Male | -0.009 | 0.037 | 0.802 |
| Side | R total (before controlling for fixed effects) | 0.217 | 0.08 | **0.002** |
| Side | R w/ fixed effects | 0.208 | 0.08 | **0.0005** |

SE: Standard error, P: P-value

**Table A2.** The consistency of individual ID for the proportion of time spent in binary position and ranked position including level of significance. Individuals’ consistency is explained by the effect of age, rank, and sex. All R values are for the random effect of individual meerkats. Binary position and ranked location both look at front vs back positions.

| Model | Predictor | Estimate | SE | P |
| --- | --- | --- | --- | --- |
| Front binary | Rank: Subordinate | -0.066 | -2.355 | **0.019** |
| Front binary | Age: Younger individuals | 0.000 | 0.024 | 0.979 |
| Front binary | Sex: Male | -0.082 | 0.036 | **0.024** |
| Front binary | Rank*Sex: Subordinate, Male | 0.087 | 0.044 | **0.047** |
| Front binary | R total (before controlling for fixed effects) | 0.117 | 0.070 | **0.013** |
| Front binary | R with fixed effects | 0.088 | 0.070 | 0.157 |
| Ranked position | Rank: Subordinate | -0.118 | 0.551 | **0.032** |
| Ranked position | Age: Younger individuals | -0.006 | 0.040 | 0.877 |
| Ranked position | Sex: Male | -0.120 | 0.062 | 0.054 |
| Ranked position | Rank*Sex: Subordinate, Male | 0.125 | 0.075 | 0.095 |
| Ranked position | R total (before controlling for fixed effects) | 0.106 | 0.064 | **0.024** |
| Ranked position | R with fixed effects | 0.087 | 0.067 | 0.159 |

**Table A3.** Estimated marginal slopes for the effect of spatial position on weight gain standardized against individual ID Δweight-Z.

| Model | Status |  | Coefficient | SE | P |
| --- | --- | --- | --- | --- | --- |
| Front | Dominant | F | -1.994 | 2.43 | 0.427 |
| Front | Dominant | M | 0.21 | 2.90 | 0.944 |
| Front | Subordinate | Older individual | -0.01 | 1.88 | 0.995 |
| Front | Subordinate | Younger individual | -2.50 | 2.53 | 0.325 |
| Back | Dominant | F | 0.84 | 1.82 | 0.645 |
| Back | Dominant | M | 0.84 | 1.82 | 0.645 |
| Back | Subordinate | Older individual | -0.01 | 1.88 | 0.995 |
| Back | Subordinate | Younger individual | -2.50 | 2.53 | 0.325 |
| Centre | Dominant | F | -1.56 | 1.40 | 0.267 |
| Centre | Dominant | M | -1.88 | 1.43 | 0.192 |
| Centre | Subordinate | Older individual | -0.01 | 1.88 | 0.995 |
| Centre | Subordinate | Younger individual | -2.50 | 2.53 | 0.325 |
| Side | Dominant | F | 2.71 | 1.99 | 0.176 |
| Side | Dominant | M | 2.81 | 1.83 | 0.126 |
| Side | Subordinate | Older individual | -0.84 | 1.22 | 0.493 |
| Side | Subordinate | Younger individual | 4.10 | 1.61 | **0.011** |

SE: Standard error, P: P-value

**Table A4.** Estimated marginal slopes for the effect of other spatial position methods on weight gain standardized against individual ID Δweight-Z.

| Model | Status |  | Coefficient | SE | P |
| --- | --- | --- | --- | --- | --- |
| Binary position | Dominant | F | -2.89 | 1.34 | **0.025** |
| Binary position | Dominant | M | -1.51 | 1.07 | 0.271 |
| Binary position | Subordinate | Older individual | 1.15 | 0.91 | 0.206 |
| Binary position | Subordinate | Younger individual | -2.12 | 1.22 | 0.084 |
| Ranked location | Dominant | F | -1.78 | 0.77 | **0.022** |
| Ranked location | Dominant | M | -0.40 | 0.77 | 0.607 |
| Ranked location | Subordinate | Older individual | 0.48 | 0.54 | 0.379 |
| Ranked location | Subordinate | Younger individual | -0.91 | 0.75 | 0.226 |

SE: Standard error, P: P-value

**Table A5.** Table for the estimated marginal slopes for the effect of spatial position on weight gain standardized against individual ID and date Δweight-Z2.

| Model | Status |  | Coefficient | SE | P |
| --- | --- | --- | --- | --- | --- |
| Front | Dominant | F | -1.67 | 2.28 | 0.466 |
| Front | Dominant | M | 0.56 | 2.72 | 0.452 |
| Front | Subordinate | Older individual | 1.12 | 1.76 | 0.527 |
| Front | Subordinate | Younger individual | -1.77 | 2.37 | 0.458 |
| Back | Dominant | F | 1.45 | 2.04 | 0.478 |
| Back | Dominant | M | 0.68 | 1.73 | 0.812 |
| Back | Subordinate | Older individual | -1.02 | 1.37 | 0.456 |
| Back | Subordinate | Younger individual | 1.04 | 1.85 | 0.576 |
| Centre | Dominant | F | -1.15 | 1.32 | 0.384 |
| Centre | Dominant | M | -2.02 | 1.35 | 0.135 |
| Centre | Subordinate | Older individual | 0.33 | 0.77 | 0.663 |
| Centre | Subordinate | Younger individual | -1.38 | 1.16 | 0.234 |
| Side | Dominant | F | 1.74 | 1.89 | 0.357 |
| Side | Dominant | M | 2.96 | 1.73 | 0.089 |
| Side | Subordinate | Older individual | -0.29 | 1.16 | 0.799 |
| Side | Subordinate | Younger individual | 2.61 | 1.52 | 0.087 |

SE: Standard error, P: P-value

**Table A6.**
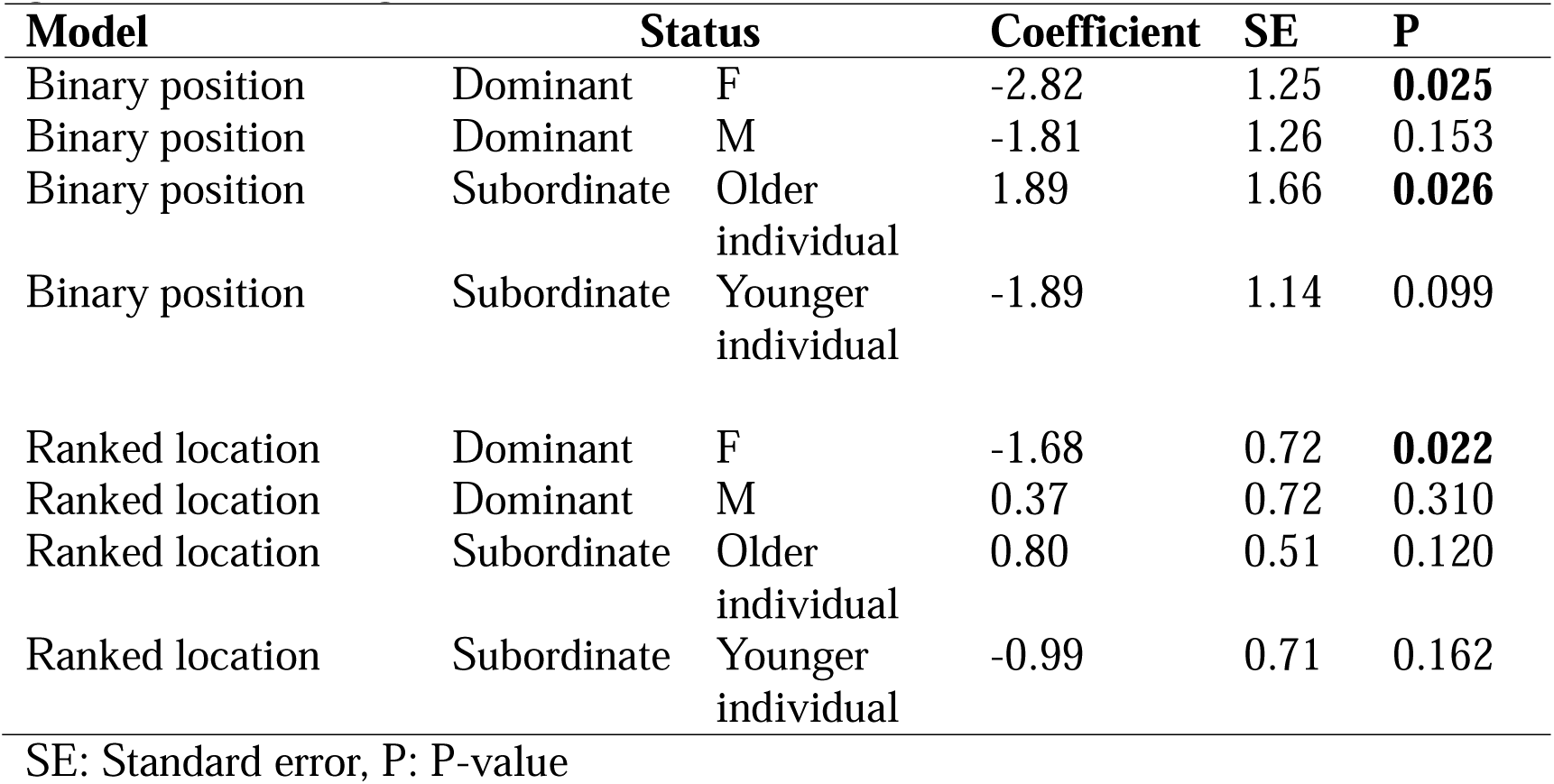
Estimated marginal slopes for the effect of other spatial position methods on weight gain standardized against individual ID and date.

| Model | Status |  | Coefficient | SE | P |
| --- | --- | --- | --- | --- | --- |
| Binary position | Dominant | F | -2.82 | 1.25 | **0.025** |
| Binary position | Dominant | M | -1.81 | 1.26 | 0.153 |
| Binary position | Subordinate | Older individual | 1.89 | 1.66 | **0.026** |
| Binary position | Subordinate | Younger individual | -1.89 | 1.14 | 0.099 |
| Ranked location | Dominant | F | -1.68 | 0.72 | **0.022** |
| Ranked location | Dominant | M | 0.37 | 0.72 | 0.310 |
| Ranked location | Subordinate | Older individual | 0.80 | 0.51 | 0.120 |
| Ranked location | Subordinate | Younger individual | -0.99 | 0.71 | 0.162 |

SE: Standard error, P: P-value

**Figure A1.**
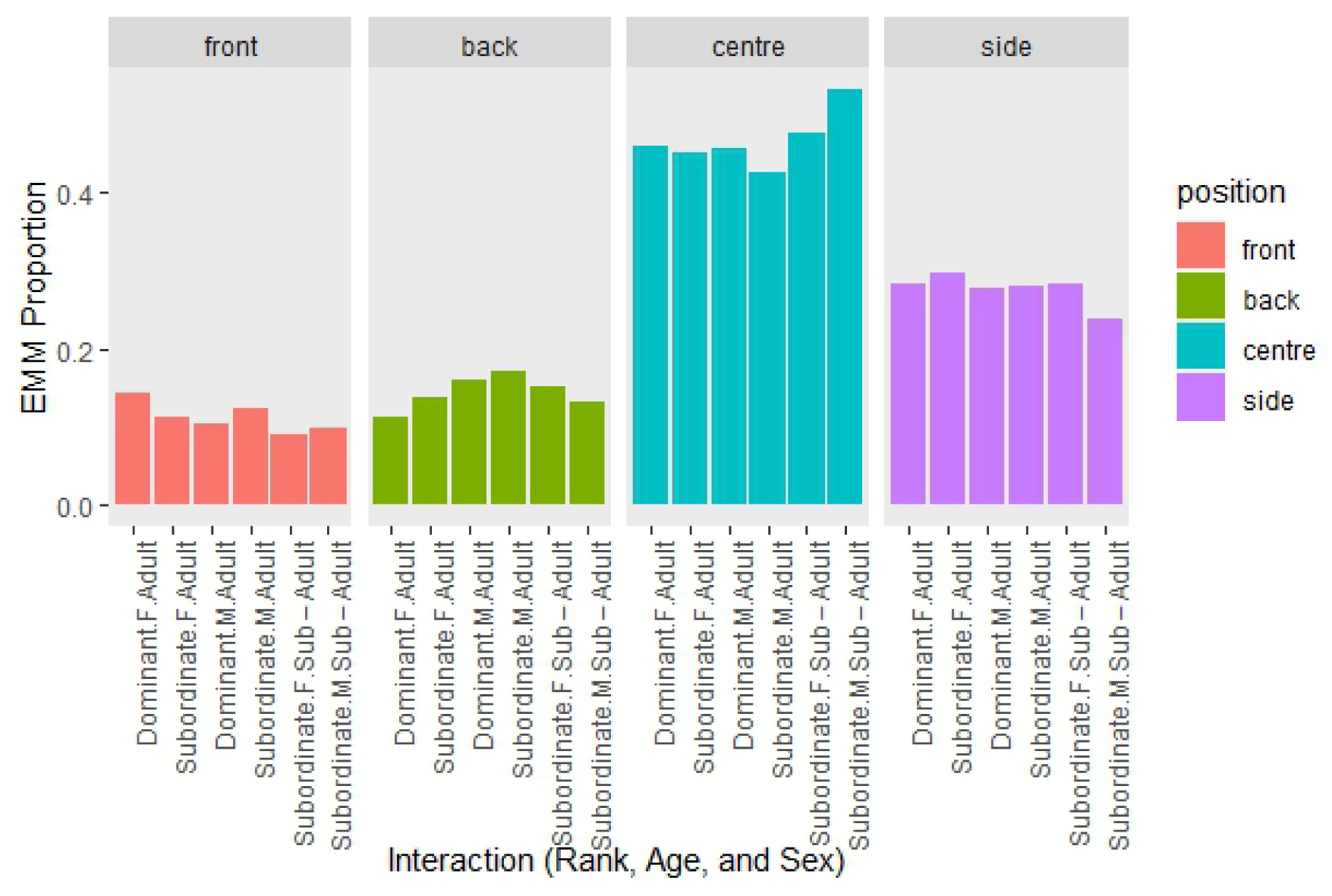
Estimated Marginal Means (EMM) of the time spent in relative position by meerkats (interaction: rank, age, and sex).

